# The Trithorax group protein dMLL3/4 instructs the assembly of the zygotic genome at fertilization

**DOI:** 10.1101/242008

**Authors:** Pedro Prudêncio, Leonardo G. Guilgur, João Sobral, Jörg D. Becker, Rui Gonçalo Martinho, Paulo Navarro-Costa

## Abstract

The transition from fertilized oocyte to totipotent embryo relies on maternally-provided factors that are synthetized and accumulated in developing oocytes. Yet, it is still unclear how oocytes regulate the expression of these embryo fate-promoting genes within the general transcriptional program of oogenesis. Here we report that the *Drosophila* Trithorax group protein MLL3/4 (dMLL3/4, also known as Trr) is essential for the transition to embryo fate at fertilization. In the absence of dMLL3/4, oocytes develop normally but fail to initiate the embryo mitotic divisions after fertilization. This incapability results from defects in both paternal genome reprogramming and maternal meiotic completion. We show that, during oogenesis, dMLL3/4 promotes the expression of a functionally coherent gene subset that is later required for the correct assembly of the zygotic genome. Accordingly, we identify the evolutionarily-conserved IDGF4 glycoprotein (known as oviductin in mammals) as a new oocyte-to-embryo transition gene under dMLL3/4 transcriptional control. Based on these observations, we propose that dMLL3/4 plays an instructive role in the oocyte-to-embryo transition that is functionally uncoupled from the requirements of normal oocyte differentiation.

## INTRODUCTION

Fertilization triggers the conversion of two terminally-differentiated haploid cells (the sperm and the oocyte) into a diploid totipotent embryo (the zygote). This transition requires a profound reprogramming of many of the cellular pathways operating in the fertilized oocyte, particularly those impinging on chromatin remodeling, cell cycle control and gene expression regulation [1]. Collectively, these changes are responsible for the oocyte-to-embryo transition, arguably one of the most dramatic cell fate changes in development.

A central feature of the oocyte-to-embryo transition is the assembly of the zygotic genome. To do so, the fertilized oocyte must merge the sperm-borne paternal genome with its maternal counterpart. Yet, this assembly is hindered by the extremely asymmetric state of parental chromatin at fertilization: the compact paternal chromatin is, in most animal species, histone-depleted and post-meiotic, while maternal chromatin remains histone-associated and meiotically-arrested [2,3]. Such asymmetry is resolved by the remodeling of the parental genomes at fertilization [4,5]. In this process, the paternal genome is decondensed through the replacement of the sperm nuclear basic proteins with maternal histones, while the maternal genome completes meiosis. Each of these events leads to the formation of a pronucleus: a nucleosome-based interphasic nucleus containing either the sperm or the oocyte-derived genomic information. The establishment of the functionally-equivalent paternal and maternal pronuclei allows the merging, at the first mitotic division, of the parental chromatin into a single zygotic genome.

A remarkable aspect of the oocyte-to-embryo transition is that it occurs in the absence of transcription [6]. Such constrain implies that the molecular determinants driving the acquisition of embryo fate, such as those involved in the assembly of the zygotic genome, must be synthetized during oogenesis and stored in the mature oocyte [7,8]. The genes coding for this unique program are referred to as maternal effect genes: a highly specialized repertoire responsible for priming and sustaining embryo development until the activation of zygotic transcription at the midblastula transition. It can be argued that the expression of these maternal effect genes during oogenesis poses a peculiar challenge to the oocyte. Indeed, developing oocytes must successfully coordinate two gene expression programs with essentially opposing outcomes: one that maintains oocyte fate as the female gamete differentiates, the other responsible for erasing such fate after fertilization.

Recently, the two chromatin regulators Mixed-lineage leukemia 3 and Mixed-lineage leukemia 4 (collectively referred to as MLL3/4) were shown to control mammalian cell fate transition to the pluripotent state during reprogramming [9]. However, despite being essential for cell fate transition, MLL3/4 were intriguingly dispensable for maintaining cell identity. MLL3/4 are part of the Trithorax group (TrxG) of proteins, a family of chromatin regulators previously proposed to function as maternal effect genes [10]. Based on these observations, we set out to determine the potential role of MLL3/4 in the oocyte-to-embryo transition.

Here, we show that *Drosophila* MLL3/4 (dMLL3/4, also known as Trr) is essential for the transition to embryo fate at fertilization but not for the maintenance of oocyte identity. More specifically, dMLL3/4 is dispensable for normal oocyte differentiation but critically required for the correct assembly of the zygotic genome at fertilization. An important aspect of such requirement is the dMLL3/4-mediated regulation of gene expression during oogenesis. Accordingly, we report the identification of a novel oocyte-to-embryo transition gene under dMLL3/4 transcriptional control.

## RESULTS AND DISCUSSION

### dMLL3/4 is essential for entry into embryogenesis

dMLL3/4 is an essential gene responsible for the monomethylation of histone H3 lysine 4 (H3K4) at enhancers and for the regulated activation of gene expression during development [11-13]. The functions of dMLL3/4 have been evolutionarily conserved [14,15], and this gene has two partially redundant mammalian homologs (MLL3 and MLL4) [16,17], both jointly required for cell fate transition but not for cell identity maintenance [9].

To test the hypothesis that dMLL3/4 promotes the oocyte-to-embryo transition at fertilization, we specifically depleted this chromatin regulator during oogenesis. For this, both an *in vivo* RNA interference (RNAi) approach and germ line mutant clone analysis (induced by the FLP/FRT recombination system) were used. The first approach ensures the post-transcriptional silencing of dMLL3/4 specifically in developing germ cells, while the second induces, in the female germ line, the homozygous mutant state of a previously identified dMLL3/4 null allele *(trr^1^*, here referred to as *dmll3/4^-/-^)* [18]. Both approaches resulted in an equally strong depletion of dMLL3/4 in whole ovaries and in early embryos (**Fig. 1A**). This result confirmed that our experimental conditions are associated with a significant reduction of dMLL3/4 levels in the female germ line.

**Figure 1.**
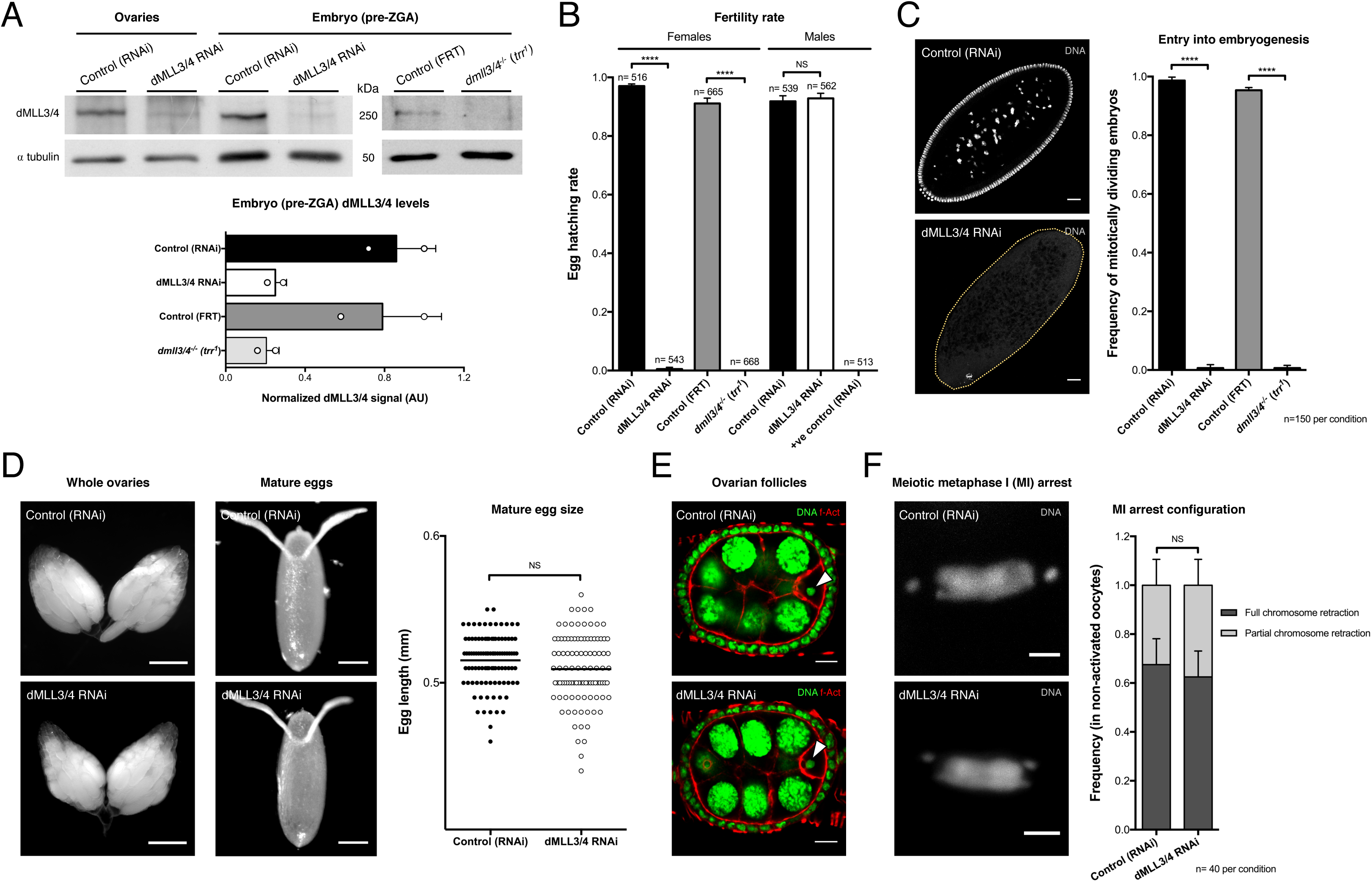
dMLL3/4 is essential for entry into embryogenesis but dispensable for oogenesis. **A**. dMLL3/4 protein levels are strongly reduced both under germ line-specific RNAi (nos-GAL4; UASp-dMLL3/4^RNAi^) and in a *dml34^-/-^* mutant *(trr^1^)*. ZGA: zygotic genome activation. The ratio between the dMLL3/4 and a-tubulin signals are expressed in arbitrary units (AU). The results of each independent experiment are plotted, bars specify mean values and horizontal lines the standard deviation. **B**. dMLL3/4 is essential for female, but not male, fertility. Error bars represent standard deviation and asterisks indicate significant difference (unpaired t-test; *P* < 0.0001; NS: no significant difference). Male germ line driver: bam-GAL4; +ve spermatogenesis control: UASp-Prp19^RNAi^. **C**. dMLL3/4-depleted eggs fail to initiate embryogenesis. The dashed yellow line delimits the egg’s cytoplasm. Unpaired t-test; *P* < 0.0001. Scale bar: 10 μm. **D**. dMLL3/4 is dispensable for morphologically normal female gonads and gametes. Mature egg size is defined by the length of its main axis and is expressed in millimetres (mm). Quantification of the egg size control group has already been published [30]. Horizontal lines specify mean values. MannWhitney U test; NS. Scale bars: 500 and 125 μm (ovaries and eggs, respectively). **E**. dMLL3/4 is dispensable for normal ovarian follicle development. Arrowheads point to the oocyte nucleus. f-Act: filamentous actin. Scale bar: 30 μm. See **Supplementary Fig. 1** for the cytological analysis of additional oogenesis stages. **F**. dMLL3/4 is dispensable for a normal meiotic metaphase I (MI) arrest. Two different configurations were observed: a tightly packaged chromosome mass (full chromosome retraction) and a more distended plate (partial chromosome retraction, as in the depicted micrographs). ANOVA; NS. Scale bar: 2 μm. Quantification of the control group has already been published [30]. See **Supplementary Fig. 2** for oocyte chromatin architecture during prophase I.

We next tested the effect of dMLL3/4 depletion on female fertility. We observed that, after being mated with wild-type males, less than 1% of all eggs laid by dMLL3/4-depleted females could hatch (n= 543 and 668, for the RNAi and *dmll3/4^-/-^* groups, respectively; **Fig. 1B**). This phenotype was limited to the female germ line, as the knockdown of dMLL3/4 during sperm differentiation had no obvious effects on male fertility (as assessed by mating dMLL3/4-depleted males with wild-type females). The origin of the female infertility phenotype could be traced to the fact that dMLL3/4-depleted eggs fail to enter embryogenesis (**Fig. 1C**). Such defect was manifested by an incapability of initiating the embryo mitotic divisions (n= 150 for each tested group).

### dMLL3/4 is dispensable for oogenesis

The germ line-specific depletion of dMLL3/4 had no obvious impact on oocyte development. Indeed, ovary morphology and mature egg size were indistinguishable between control and dMLL3/4-depleted conditions (**Fig. 1D**). Supporting this observation, we failed to identify any cytological defects in either the germ cell or somatic compartments of depleted ovaries (**Fig. 1E** and **Supplementary Fig.1**). Equally unaffected was the meiotic metaphase I (MI) arrest of mature, non-activated eggs. More specifically, dMLL3/4 depletion did not disturb the retraction of the meiotic chromosomes into the MI plate (n= 40, **Fig. 1F**), nor did it impact prophase I oocyte chromatin architecture (n= 10, **Fig. 1E** and **Supplementary Fig.2**).

Collectively, the germ line-specific depletion of dMLL3/4 did not result in any noticeable disturbances in oogenesis, despite critically impairing the entry into embryogenesis. The fact that dMLL3/4 is dispensable for morphologically normal oogenesis contrasts with previous observations for other SET domain histone methyltransferases across different species [19-22].

### dMLL3/4 is required for the reprogramming of the paternal genome at fertilization

Why do dMLL3/4-depleted eggs fail to enter embryogenesis? We observed that the depletion of this chromatin remodeler during oogenesis blocks the reprogramming of the paternal genome after fertilization (**Fig. 2A**). More specifically, after being mated with wild-type males, more than 90% of all eggs laid by dMLL3/4-depleted females are unable to convert the tightly compacted sperm-borne paternal genome into the decondensed chromatin of the paternal pronucleus (PN; n= 36 and 51, for the RNAi and *dmll3/4^-/-^* groups, respectively). Indeed, in the absence of dMLL3/4, the sperm genome retains its compact needle-like shape after fertilization, whereas this configuration is rapidly converted into a round decondensed structure in controls (**Fig. 2A**). Given the extremely fast kinetics of *Drosophila* sperm decondensation (in normal conditions the first zygotic division occurs just 15 minutes after sperm entry) [23], the previously reported *sra^-/-^* mutant *(sra^A108^/sra^A426^)* was used as control [24]. In this egg activation mutant, the reprogramming of the sperm-borne genome is blocked immediately after decondensation. Consequently, in fertilized eggs from this genetic background, the paternal genome remains arrested in the shape of the decondensed male PN (**Fig. 2A**).

**Figure 2.**
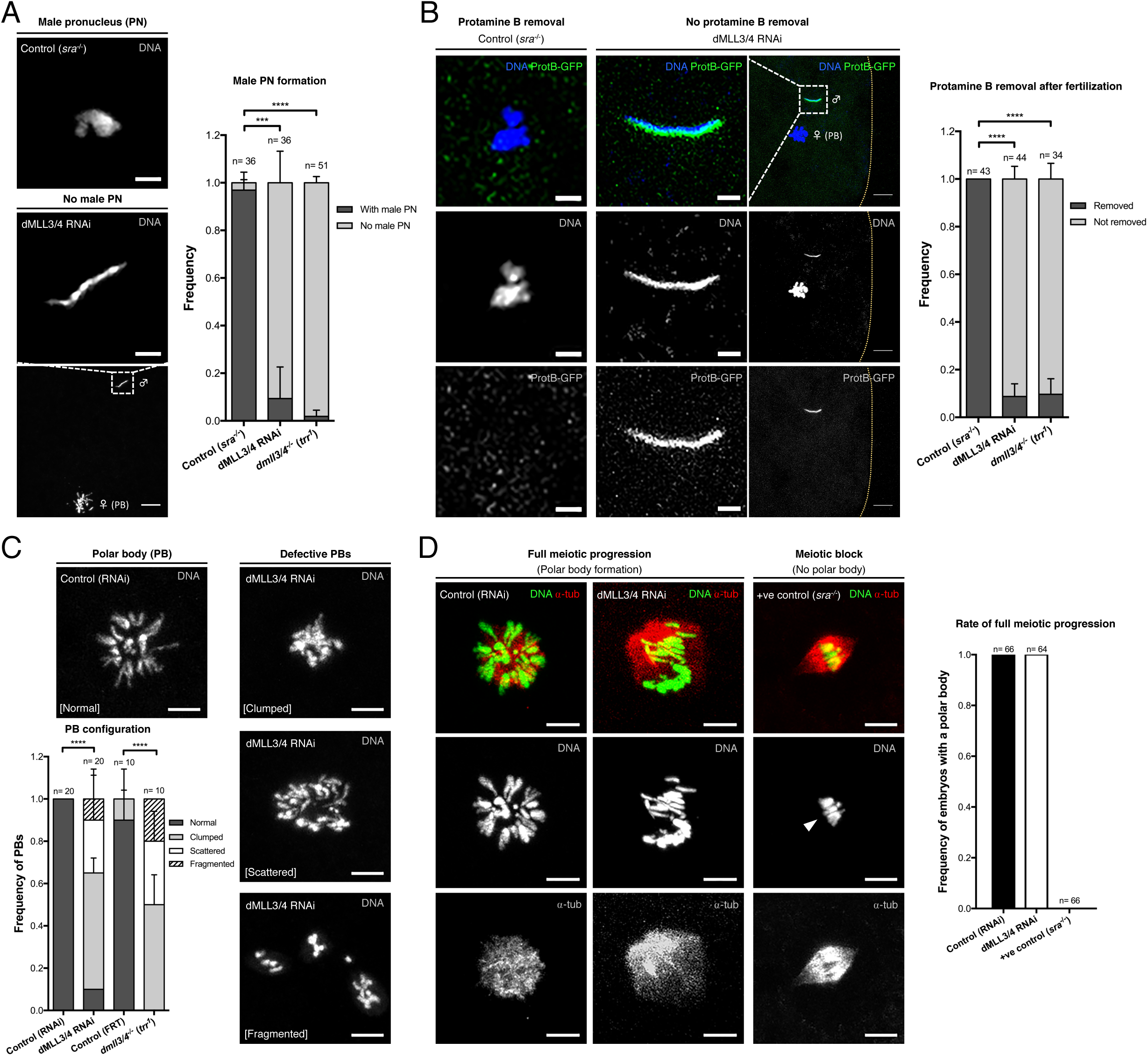
dMLL3/4 is critically required for the remodeling of the two parental genomes at fertilization. **A**. dMLL3/4-depleted eggs are incapable of forming the male pronucleus (PN) after fertilization. In such conditions, paternal chromatin (♂) remains in a condensed sperm-like state. The *sra^-/-^* mutant was used as control for male PN formation. ♀: female meiotic products (polar body: PB). Error bars represent standard deviation and asterisks indicate significant difference (ANOVA; *P ≤* 0.0002). Scale bars: 2 μm (for ♂ insets) and 10 μm. **B**. dMLL3/4-depleted eggs are unable to remove protamine B from the sperm DNA. ANOVA; *P* < 0.0001. The dashed yellow line delimits the egg’s cytoplasm. Scale bars: 2 μm (for ♂ insets) and 10 μm. **C**. dMLL3/4 is required for normal female meiotic completion. PB chromatin morphology was used as read-out for successful meiotic completion. dMLL3/4 depletion led to significant deviations to the normal PB rosette conformation. ANOVA; *P* < 0.0001. Scale bars: 5 μm. **D**. Meiotic progression is not affected in dMLL3/4-depleted eggs. PB formation was used as read-out for the progression through the two meiotic divisions. The *sra^-/-^* mutant was used as positive control for a female meiotic block. Arrowhead: meiosis I-arrested chromatin (compare size with that of PBs). Scale bars: 5 μm.

To understand why the sperm-borne paternal genome fails to be decondensed in dMLL3/4-depleted eggs, RNAi females were mated with males carrying endogenously fluorescent protamine B (ProtB-GFP) [25]. Protamine B is one of the major nuclear proteins incorporated into sperm DNA during spermatogenesis, and the formation of the decondensed paternal PN after fertilization requires its active removal (alongside other basic nuclear proteins) from the sperm-borne genome [26]. We observed that the depletion of dMLL3/4 during oogenesis resulted in eggs incapable of ensuring protamine removal after fertilization (**Fig. 2B**). Indeed, in more than 90% of the fertilized eggs laid by dMLL3/4-depleted females, paternal chromatin retained the ProtB-GFP fluorescent signal (n= 44 and 34, for the RNAi and *dmll3/4^-/-^*groups, respectively). In contrast, no ProtB-GFP signal was ever recorded in the control *(sra^-/-^* mutant; n= 43; **Fig. 2B**). Similar to our experiments with wild-type sperm, the genome of ProtB-GFP sperm also remained in a compact configuration in dMLL3/4-depleted eggs, while it was extensively decondensed in controls.

Based on these observations, we conclude that the expression of dMLL3/4 during oogenesis confers oocytes the ability to decondense the paternal genome after fertilization. The functional basis of this ability resides in the removal of DNA-compacting nuclear proteins from the sperm-borne chromatin.

### dMLL3/4 is required for maternal meiotic completion

We found that the role of dMLL3/4 in post-fertilization development went beyond the decondensation of the paternal genome. In fact, dMLL3/4 was also required for the successful completion of female meiosis. Meiotic completion is an essential aspect of the oocyte-to-embryo transition: it erases the cell cycle stage asymmetry between the maternal and paternal genomes at fertilization. A particularity of female meiosis is that out of the four resulting haploid nuclei, only one will contribute to the next generation [27]. All three remaining nuclei (known as polar bodies – PBs) eventually degenerate. In *Drosophila*, these three nuclei fuse, after DNA replication, in the cytoplasm of the egg [28,29]. The condensed chromosomes of this single PB organize into a characteristic rosette shape, commonly used as readout for successful meiotic completion (**Fig. 2C**) [30]. dMLL3/4 depletion critically impaired the formation of a normal PB rosette. This PB configuration - indicative of normal meiotic completion - was only detected in less than 10% of all dMLL3/4-depleted eggs (n= 20 and 10, for the RNAi and *dmll3/4^-/-^*groups, respectively). In all other cases, PB chromatin was severely affected: chromosomes often collapsed into an indistinct chromatin mass (clumping), failed to properly assemble into a rosette (scattering), or were outright dispersed in the cytoplasm of the egg (fragmentation; **Fig. 2C**). Despite these obvious meiotic defects, dMLL3/4-depleted eggs invariably completed meiosis (**Fig. 2D**). In fact, the formation of structurally abnormal PBs in all dMLL3/4-depleted eggs confirmed their capability of progressing, upon egg activation, from the meiotic metaphase arrest to the post-meiotic state (n= 64 and 66, for the RNAi and control groups, respectively). This observation stands in stark contrast with egg activation mutants, such as *sra^-/-^*, which are incapable of progressing from the meiotic metaphase arrest (**Fig. 2D**) [24,31].

Collectively, these observations indicate that the expression of dMLL3/4 during oogenesis does not interfere with meiotic cell cycle progression but is essential for the successful completion of female meiosis.

### dMLL3/4 regulates the expression of a functionally coherent gene subset

Having established that dMLL3/4 is required for the correct remodeling of both the paternal and maternal genomes at fertilization, we next set out to characterize the mechanistic basis of this requirement. dMLL3/4 is a transcriptional activator belonging to the TrxG protein family [14,15]. TrxG complexes promote the expression of developmental genes across multiple cellular contexts. Therefore, we hypothesized that dMLL3/4 activates, during oogenesis, the expression of genes that will be later required for post-fertilization development. To test this hypothesis, we performed a transcriptomic analysis of dMLL3/4-depleted eggs. Such analysis revealed that the depletion of dMLL3/4 during oogenesis had a minor effect on the transcriptome: only 1.5% of the genes were differentially expressed (212 out of 14401; adjusted p-value <0.05; **Fig. 3A**). Consistent with the role of dMLL3/4 as a transcriptional activator, the majority (69%) of the differently expressed genes were downregulated (146 downregulated genes, excluding dMLL3/4, *vs*. 65 upregulated; **Supplementary Table 1**). The subtle transcriptomic imbalance of dMLL3/4-depleted eggs was associated with an equally moderate impact on total H3K4 methylation levels in the ovary (**Supplementary Fig. 3**).

**Figure 3.**
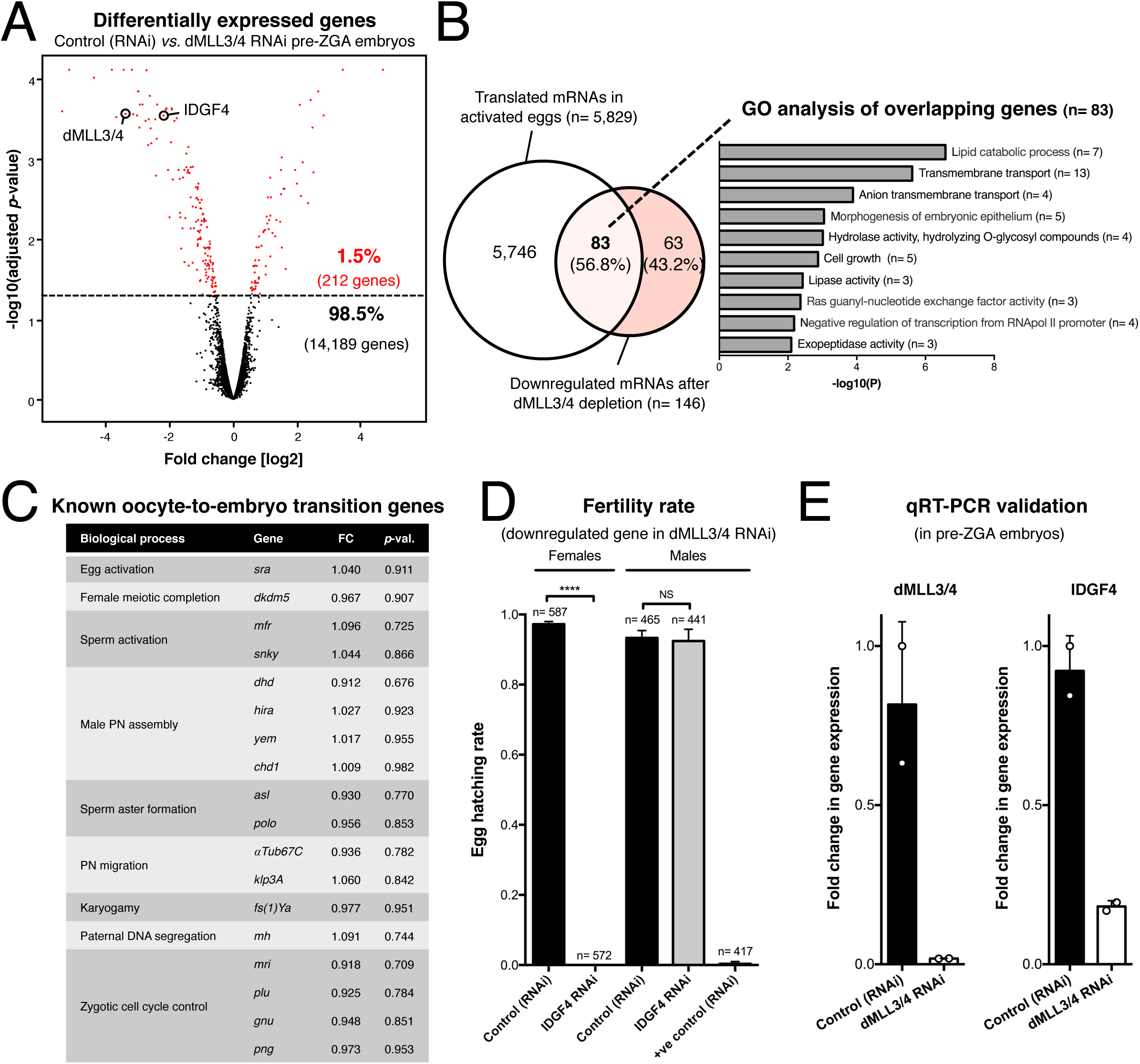
dMLL3/4 defines the expression of a small, functionally coherent maternal gene subset. **A**. Germ line-specific depletion of dMLL3/4 during oogenesis affects a small number of maternal transcripts. Differently expressed genes are represented in red (adjusted p-value cut-off: 0.05). All embryos were manually isolated prior to the onset of zygotic genome activation (pre-ZGA). See panel **E** for qRT-PCR validation of selected genes and **Supplementary Table 1** for expression values. **B**. The majority of dMLL3/4-activated genes are preferentially translated during egg activation. Gene ontology (GO) analysis of this gene subset identified embryogenesis-related GO terms. Translated mRNAs in activated eggs were extracted from a previously published ribosome footprinting dataset [32]. **C**. The expression of known oocyte-to-embryo transition genes is not significantly altered in dMLL3/4-depleted eggs. FC: linear fold change in dMLL3/4 RNAi; *p*-val.: adjusted *p*-value; PN: pronucleus. See **Supplementary Table 2** for full expression values. **D**. IDGF4 is essential for female, but not male, fertility. See **Fig. 4** for the role of IDGF4 in the oocyte-to-embryo transition. Error bars represent standard deviation and asterisks indicate significant difference (unpaired t-test; *P* < 0.0001; NS: no significant difference). Male germ line driver: bam-GAL4; +ve spermatogenesis control: UASp-Prp19^RNAi^. **E**. Real-time quantitative reverse transcription PCR (qRT-PCR) validates the downregulation observed in microarray data. Expression levels were normalized against a highly expressed reference gene (RpL32).

Further analysis of the dMLL3/4-defined gene expression program revealed three noteworthy observations. First, the majority (57%) of the dMLL3/4-activated genes are, under normal conditions, translated at the oocyte-to-embryo transition. More specifically, by comparing our transcriptomic data with a previously published ribosome footprinting dataset of activated eggs [32], we found that 83 out of the 146 downregulated genes were translated at this transition stage (**Fig. 3B**; **Supplementary Table 1**). Secondly, consistent with the observed phenotypes, oogenesis-related genes are conspicuously absent from the dMLL3/4-activated gene expression program (**Fig. 3B**). Accordingly, the gene ontology (GO) analysis of the 83 dMLL3/4-activated genes translated at the oocyte-to-embryo transition failed to detect any enrichment for oogenesis-related processes (such as “female gamete generation” or “female meiosis”) [33]. On the other hand, embryogenesis-related GO terms (such as “morphogenesis of embryonic epithelium” and “negative regulation of transcription from RNApol II promoter”) stood out from the list of enriched terms. Thirdly, dMLL3/4 does not regulate the expression of any gene previously associated with the assembly of the zygotic genome [4,26,30]. In particular, all known *Drosophila* oocyte-to-embryo transition genes were expressed at levels similar to controls (**Fig. 3C** and **Supplementary Table 2**).

Based on these data, we conclude that dMLL3/4 promotes, during oogenesis, the expression of a small subset of genes preferentially translated during the acquisition of embryo fate. Could the deregulation of any of them be responsible for the incapability of dMLL3/4-depleted eggs to initiate embryogenesis?

### IDGF4 is a new, dMLL3/4-regulated, oocyte-to-embryo transition gene

To address the functional significance of the transcriptomic defects of dMLL3/4-depleted eggs, we individually silenced, during oogenesis, the more severely downregulated genes (**Supplementary Table 3**). By analysing how this silencing affected female fertility, we identified a new oocyte-to-embryo transition gene: Imaginal disc growth factor 4 (IDGF4) [34].

The female germ line-specific depletion of IDGF4 had a critical impact on fertility: after mating with wild-type males, none of the eggs laid by IDGF4-depleted females hatched (n= 572 and 587, for the RNAi and control groups, respectively; **Fig. 3D**). The reduced expression of IDGF4 in dMLL3/4 RNAi eggs was confirmed by real-time quantitative reverse transcription PCR (**Fig. 3E**), and is consistent with the fact that IDGF4 harbours a predicted Polycomb/Trithorax response element [35]. Importantly, IDGF4 depletion recapitulated several of the oocyte-to-embryo transition defects of the dMLL3/4 RNAi. More specifically, despite their normal morphology, IDGF4-depleted eggs failed to initiate the embryo mitotic divisions (n= 150 for each tested group; **Fig. 4A**, compare with **Fig. 1C**). In addition, these eggs had maternal meiotic completion defects equivalent to those observed in the dMLL3/4 RNAi. Such defects were illustrated by the clumping, scattering or outright fragmentation of PB chromosomes (n= 20 per group; **Fig. 4B**, compare with **Fig. 2C**).

**Figure 4.**
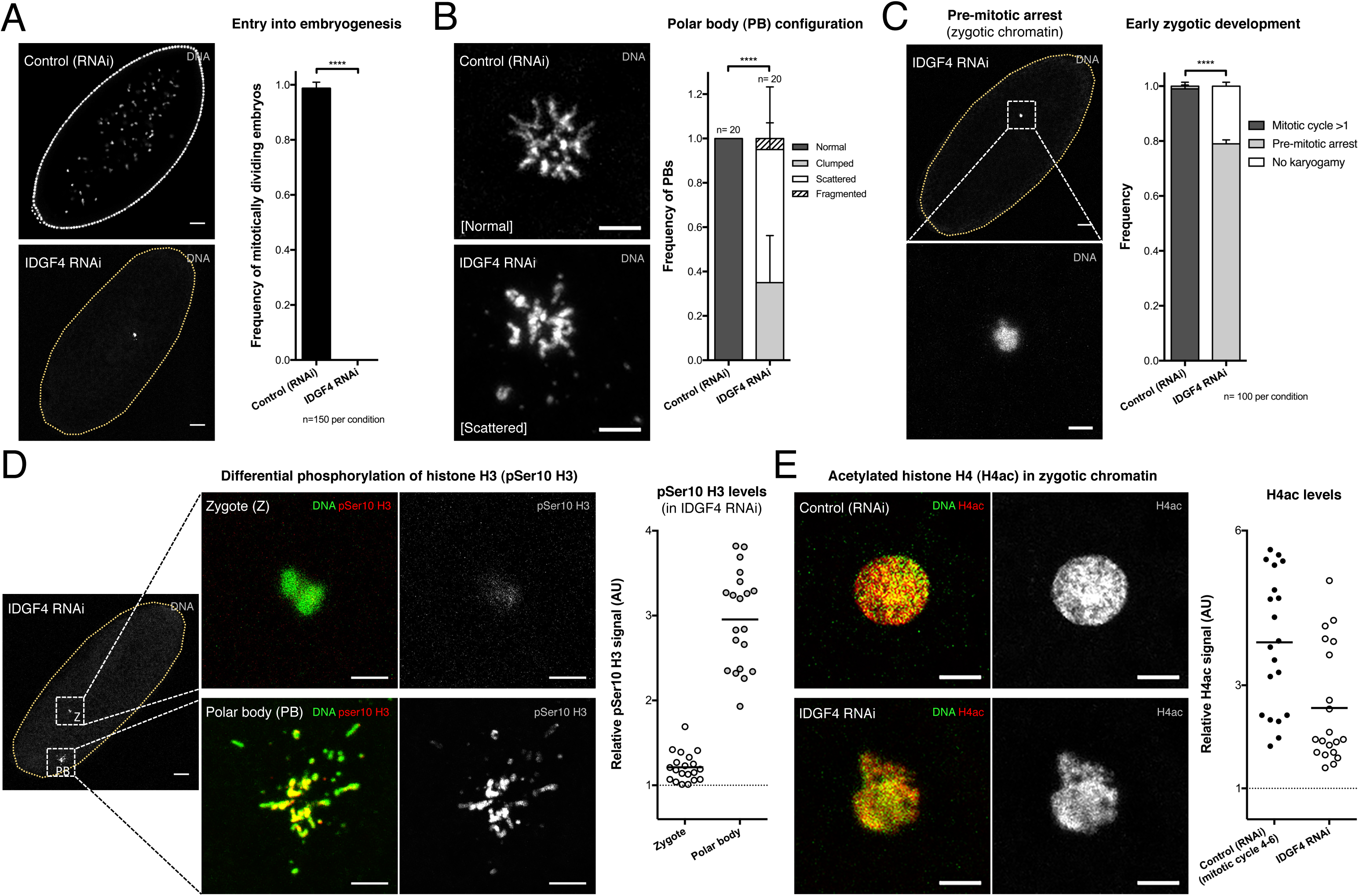
The dMLL3/4-regulated glycosyl hydrolase IDGF4 is a new oocyte-to-embryo transition gene. **A**. IDGF4-depleted eggs fail to initiate embryogenesis. Compare with the similar observation recorded in dMLL3/4-depleted eggs (see **Fig. 1C**). Driver: *nos-* GAL4. The dashed yellow line delimits the egg’s cytoplasm. Error bars represent standard deviation and asterisks indicate significant difference (unpaired t-test; *P* < 0.0001). Scale bar: 30 μm. **B**. IDGF4 is required for normal maternal meiotic completion. Polar body (PB) chromatin morphology was used as read-out for successful meiotic completion. Compare with the similar observation recorded in dMLL3/4-depleted eggs (see **Fig. 2C**). ANOVA; *P* < 0.0001. Scale bar: 5 μm. **C**. IDGF4-depleted eggs assemble mitotically-deficient zygotic chromatin. Such chromatin was arrested at a pre-mitotic state. ANOVA; *P* < 0.0001. Scale bars: 30 μm (egg) and 5 μm (inset). **D**. IDGF4 is specifically required for the phosphorylation of histone H3 in zygotic chromatin. Although histone H3 serine 10 phosphorylation (pSer10 H3) was virtually undetectable in zygotic chromatin (Z), the same embryos displayed high levels of this modification in their PB chromosomes. Signal quantification is expressed in fluorescence arbitrary units (AU). Scale bars: 30 μm (egg) and 5 μm (insets). **E**. IDGF4 depletion does not abrogate the acetylation of histone H4 (H4ac) in zygotic chromatin. For comparison, the early zygotic chromatin of control embryos (mitotic cycle 4-6) is depicted. Scale bars: 5 μm.

IDGF4 is one of the six members of the IDGF family of glycosyl hydrolases [34,36]. IDGFs are required for insulin-dependent cell proliferation, polarization and motility. Their activity has also been linked to developmental processes such as extracellular matrix formation, innate immunity, wound healing and tissue morphogenesis [37-39]. We observed that IDGF4 is essential for the transition from the meiotic to the mitotic program at fertilization. Indeed, most (78%) IDGF4-depleted eggs assembled zygotic chromatin but were incapable of initiating mitosis (n= 100 for each tested group; **Fig. 4C**). This pre-mitotic arrest was consistent with barely detectable levels of histone H3 serine 10 phosphorylation (pSer10 H3, a marker of chromosome condensation) in zygotic chromatin (n= 20 for each tested group; **Fig. 4D**) [40]. Notably, not only pSer10 H3 was abundantly detected in the polar bodies of the same embryos, but also the levels of acetylated histone H4 in zygotic chromatin were high (n= 20 for each tested group; **Fig. 4E**). Both observations strongly suggest that the low pSer10 H3 zygotic signal of IDGF4-depleted embryos specifically reflects a mitotic entry defect rather than a generalized deregulation of the post-fertilization epigenome.

In summary, these data support the notion that IDGF4 is a new, dMLL3/4-regulated, oocyte-to-embryo transition gene. Such observation validates the instructive role of dMLL3/4 in the acquisition of embryo fate.

## CONCLUSIONS

According to our model, dMLL3/4 regulates, during oocyte development, the expression of a small gene repertoire that is required for the acquisition of embryo fate (**Fig. 5**). Such regulation may have evolved to better accommodate the expression of specific maternal effect genes within the general transcriptional program of oogenesis. dMLL3/4-mediated gene expression can thus be considered an additional regulatory layer in the complex orchestration of the oocyte-to-embryo transition. Although this transition is largely dependent on the profound posttranscriptional changes taking place at egg activation [32,41,42], our observations emphasize the importance of germ cell transcriptional regulation in the acquisition of embryo fate.

**Figure 5.**
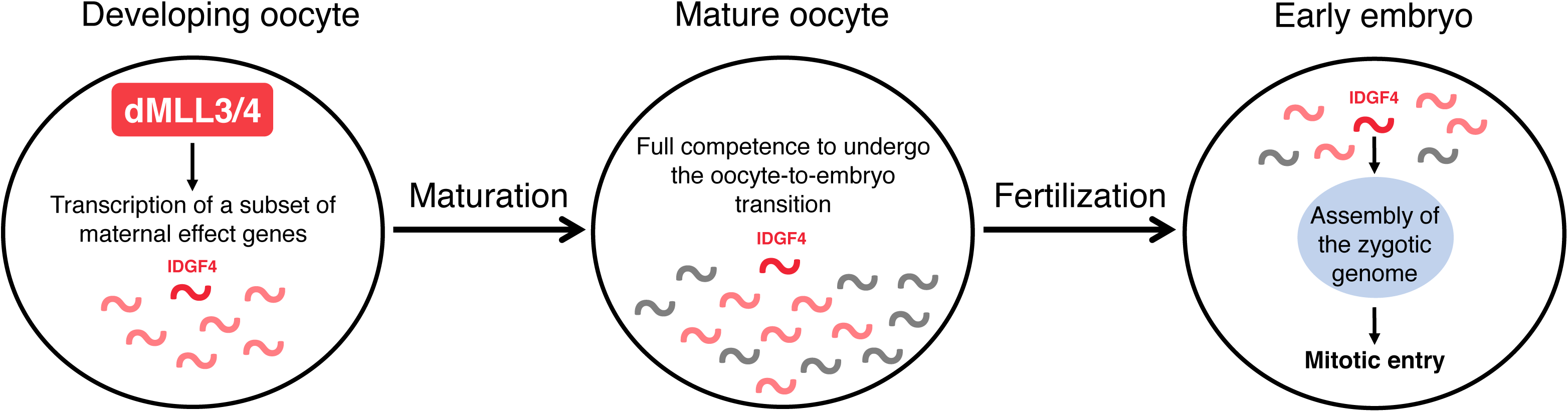
Proposed model for the dMLL3/4-regulated acquisition of embryo fate at fertilization. As oocytes develop, the Trithorax group protein dMLL3/4 promotes the expression of a small subset of genes. The products of these genes (in red) are dispensable for normal oogenesis but their storage in the mature oocyte (alongside other maternal effect genes, in grey) is fundamental for the correct assembly of the zygotic genome at fertilization. The glycosyl hydrolase IDGF4 is one of the dMLL3/4-regulated genes critically required for the oocyte-to-embryo transition.

How does dMLL3/4 regulate transcription during oogenesis? Despite containing a H3K4 methyltransferase domain, the catalytic activity of dMLL3/4 has been shown to be dispensable, in unperturbed conditions, for development, viability and fertility [13]. dMLL3/4 is part of an evolutionarily-conserved multiprotein complex (dMLL3/4-COMPASS) that localizes to enhancer regions of the genome [14,15]. We envisage that enhancers for specific maternal effect genes have a distinct chromatin environment amenable to the binding of dMLL3/4-COMPASS. The possibility that specific gene subsets harbor a defined epigenetic signature in germ cells is not without precedent. Indeed, key developmental genes have been previously shown to be maintained in a poised epigenetic state in mouse germ cells [43]. Quite fittingly, such epigenetic state is believed to serve as a priming mechanism for the acquisition of totipotency following fertilization [44].

Previous observations suggest that it is the binding at enhancers and not the catalytic activity of dMLL3/4 that is essential for the activation of gene expression. More specifically, in mouse embryonic stem cell (mESC) lines, the recruitment of RNA polymerase II and enhancer RNA synthesis are drastically reduced in the absence of MLL3 and MLL4, but marginally affected in a catalytic dead condition [45]. The dispensable nature of dMLL3/4-mediated methylation is particularly evident in the female germ line, as our results indicate that total H3K4 methylation levels in the ovary remain largely unchanged in the absence of dMLL3/4. More recently, it has been proposed, also in mESC lines, that the catalytic activity of MLL3 and MLL4 may in fact facilitate the binding of additional chromatin remodelers to enhancers [46]. Collectively, these observations illustrate the multi-layered role of dMLL3/4-COMPASS in gene expression regulation. A better understanding of such role may be of significance in the context of complex human diseases such as infertility.

Two other TrxG proteins are known to confer oocytes the capability of sustaining post-fertilization development. MLL2 is required during mammalian oogenesis for the subsequent activation of the zygotic genome [20], and we and others have shown that the depletion of dKDM5 in developing *Drosophila* oocytes impairs early embryo development [30,32]. By adding dMLL3/4 to this list, our work underlines the importance of TrxG proteins in the transition to totipotency. In this sense, modulating the expression of TrxG proteins, such as MLL3/4, during somatic cell reprogramming might increase the overall efficiency of this technique. An analogous approach targeting one of the vertebrate orthologs of dKDM5 has already been shown to improve the development of nuclear transfer embryos [47].

The relevance of our findings may even extend to assisted reproduction. We observed that the dMLL3/4-regulated glycosyl hydrolase IDGF4 has a mitogenic effect on the early embryo. The mammalian ortholog of IDGF4 is OVGP1, also known as oviductal glycoprotein 1 [36]. OVGP1 localizes to the oviduct – the site of mammalian fertilization – where it has been shown to be a component of the extracellular matrix of ovulated oocytes and early embryos [48,49]. Despite being important for fertilization and early embryogenesis across different mammalian species, the exact function of OVGP1 remains elusive [50]. Accordingly, OVGP1 has been associated with processes as wide-ranging as sperm capacitation and blastocyst formation [51-53]. Our results indicate that the activation of embryo mitotic divisions is the critical function of IDGF4/OVGP1. Given that IDGF4 has been linked to the acquisition of cellular responsiveness to signaling cues [54], this glycoprotein may be involved in the transduction of fertilization-dependent signals required by the embryo mitotic cascade. In this context, the supplementation of human *in vitro* fertilization (IVF) culture media with OVGP1 may ultimately increase the developmental potential of fertilized oocytes. Indeed, the addition of this extracellular glycoprotein to IVF media has been shown to significantly improve early embryo development rates in several livestock species [55,56]. Bridging such result to the clinical setting could represent an important advance in human assisted reproduction techniques.

## METHODS

No statistical methods were used to predetermine sample size. Ovary imaging and prophase I oocyte chromatin analysis experiments were intentionally randomized, and the investigators blinded to allocation during experiments and outcome assessment.

### *Drosophila* rearing conditions

*Drosophila melanogaster* flies were raised at 25°C in polypropylene vials (51 mm diameter) containing enriched medium (cornmeal, molasses, yeast, soya flour and beetroot syrup). For ovary analysis, flies were transferred, 24h prior to being tested, to polystyrene vials (23.5 mm diameter) containing standard medium (cornmeal, molasses, yeast and sucrose). For embryo analysis, flies were transferred 48h before testing to egg laying cages (with 60 mm diameter apple juice agar plates). In both cases, fresh yeast paste was provided. Tested flies were collected 3 to 7 days post-eclosion. All females under analysis were kept with wild type males (Oregon-R) except when they were mated with ProtB-GFP males to assess paternal genome reprogramming at fertilization.

### Germ line-specific RNAi

The Gal4-UASp system was used to silence genes of interest specifically in the female germ line [57,58]. In all reported experiments the silencing was induced from early oogenesis (germ line stem cell) to the mature egg stage, using the well-established nos-GAL4 driver [59]. dMLL3/4 was silenced using a publically available UASp-RNAi line (Bloomington stock no. 36916). For the list of UASp-RNAi lines against dMLL3/4-regulated genes, please consult **Supplementary Table 3**. As controls, the RNAi response was induced, by nos-GAL4, against a sequence not present in the genome of the tested flies (mCherry fluorophore; Bloomington stock no. 35785). For the male fertility tests, selected genes were specifically silenced in the male germ line using bam-GAL4, a well-characterized spermatogenesis driver [60].

### Maternal mutant generation

The previously published embryonic lethal *trr^1^* null allele was used to generate *dmll3/4^-/-^* maternal mutant embryos [18]. For this, the FLP/FRT Ovo^D^ recombination system was employed [61,62]. Briefly, *Ovo^D^, FRT101/Y; FLP38* males (Bloomington stock no. 1813) were crossed to *trr^1^, FRT101/FM7*females. Recombination was induced by heat shocking third instar larvae for 1 hour at 37°C. For controls, recombination was induced using *FRT101* females (Bloomington stock no. 1844).

### Additional *Drosophila* strains

Two previously published loss of function alleles affecting the *Drosophila* calcipressin gene *sarah (sra^A108^* and *sra^A426^)* were used to generate the *sra^-/-^* egg activation mutant *(sra^A108^/sra^A426^* transheterozygote) [24]. The reprogramming of the paternal genome after fertilization was tested by mating females under analysis with ProtB-GFP males carrying a fusion of Protamine B with enhanced green fluorescent protein *(Sp/CyO; ProtB-GFP)* [25]. An UASp fly stock carrying a double stranded RNA against an essential gene (the spliceosome subunit Prp19) was used as positive control for spermatogenesis phenotypes [63].

### Antibodies

The following primary antibodies were used for immunofluorescence: mouse anti-a-Tubulin (1:500 dilution, Sigma T9026); mouse anti-Orb (clones 4H8 and 6H4, 1:30 dilution each, Developmental Studies Hybridoma Bank); rabbit anti-pSer10 H3 (1:500 dilution, Upstate 06-570) and rabbit anti-H4ac (1:500 dilution, Millipore 06-866). The following primary antibodies were used for immunoblotting: rabbit anti-dMLL3/4 (1:1000 dilution, from both the Alexander Mazo and Ali Shilatifard labs) [11,12]; rabbit anti-H3K4me1 (1:5000 dilution, Active Motif 39298); rabbit anti-H3K4me2 (1:5000 dilution, Active Motif 39142); rabbit anti-H3K4me3 (1:500 dilution, Active Motif 39160); rabbit anti-H3 (1:8000 dilution, Cell Signalling Technology #9715) and mouse anti-α-Tubulin (1:50000 dilution, Sigma T6199).

Secondary detection was performed with Cy3, Cy5 and HRP-conjugated antibodies at 1:1000 (immunofluorescence) and 1:4000 (immunoblotting) dilutions (Jackson ImmunoResearch).

### Protein immunoblotting

Maternal protein extracts were prepared from pre-zygotic genome activation (ZGA) embryos as previously described [30,63]. Briefly, early embryos (less than 1 hour post-laying) were dechorionated with a 50% commercial bleach solution and manually selected for the lack of morphological hallmarks of a post-ZGA state (selection for the absence of pole cells and cortical nuclei). Maternal protein extracts were obtained by lysing the embryos with a needle in Laemmli sample buffer and heating for 5 minutes at 100°C. Fifteen embryos were selected per genotype per experiment. Ovary protein extracts were enriched for core histones using a histone purification mini kit (Active Motif). Twenty ovary pairs were included per sample.

Protein samples were run on 6% or 15% SDS-PAGE gels (for dMLL3/4 and H3K4 methylation analysis, respectively) and transferred to nitrocellulose membranes (Amersham) for one hour at 100V. Membranes were blocked for one hour at room temperature in 5% non-fat milk in PBS-T [0.1% Tween 20 in PBS (both from Sigma-Aldrich)], followed by an overnight incubation with the primary antibody (at 4°C) in 1% non-fat milk in PBS-T. After washing, membranes were blocked for 15 minutes in 5% non-fat milk in PBS-T and then incubated for 2 hours (at room temperature) with the secondary antibody in 1% non-fat milk in PBS-T. Following another round of washing, membranes were incubated with ECL solution for 1 minute. Protein detection was performed using a ECL Hyperfilm (Amersham). A minimum of two independent experiments was conducted for each experimental condition.

### Fertility tests

Fertility was tested as previously described [30]. In either male or female fertility tests, egg laying cages with 20 females and 10 males were maintained at 25°C for 48 hours prior to analysis. Tested genotypes were mated with a wild type strain (Oregon-R). All flies were 3 to 7 days post-eclosion and analyses were performed on two consecutive days. Laid eggs were collected for 30 minutes in apple juice agar plates and further incubated for 48 hours at 25°C. The total number of eggs and the number of hatched eggs were recorded upon collection and 48 hours afterwards, respectively. Fertility rate was determined as the number of hatched eggs divided by the total number of eggs. Four independent experiments were conducted for each experimental condition.

### Embryo imaging

Egg laying cages with 40 females and approximately 20 males were maintained at 25°C for 48 hours prior to analysis. Embryos were collected in apple juice agar plates (up to 30 minutes after egg laying) and immediately processed. Collection times went up to 1 hour after egg laying followed by 2 additional hours of incubation at 25°C when assessing entry into embryogenesis. Embryo processing started with dechorionation in a 50% commercial bleach solution. Embryos were then fixed and devitellinized in a 1:1 heptane-methanol mix for 5 minutes with shaking. Following rehydration, embryos were stained as previously described [64]. Briefly, embryos were blocked for 1 hour in BBT [PBS-T supplemented with 1% (w/v) bovine serum albumin and 1% (w/v) donkey serum (both from Sigma-Aldrich)]. Primary antibody incubation was performed overnight at 4°C in BBT. After washing, embryos were incubated in the secondary antibody solution (in BBT, for 1 hour at room temperature). DNA was stained either with SYTOX Green (Invitrogen) or Hoechst 33342 (Thermo Fisher). For SYTOX Green, embryos were incubated for 30 minutes in a 1:5000 dilution in PBS-T supplemented with 5 μg/ml RNase A (Sigma-Aldrich). For Hoechst 33342, the incubation time for a 5 μg/ml solution in PBS-T was 10 minutes. Embryos were mounted in fluorescence mounting medium (Dako) and images were acquired with either a 40x HCX PL APO CS oil immersion objective (numerical aperture: 1.25-0.75) or a 63x HCX PL APO oil immersion objective (numerical aperture: 1.40-0.60) on a Leica SP5 confocal microscope. Four independent experiments were conducted for each experimental condition. For the measurement of mature eggs, these were photographed (after a one hour collection) using a camera-coupled Leica MZ12.5 stereomicroscope (Plan APO 1.0x objective, amplification: 64x). Egg size corresponds to the length of its main axis, as measured using the ImageJ software (v1.48i; National Institutes of Health). Since this experiment (**Fig. 1D**) was part of a larger dataset, the quantification of the control group has already been published [30].

### Ovary imaging

Adult ovaries (10 ovary pairs per sample per experiment) were processed as previously described [30]. Briefly, ovaries were dissected in PBS and fixed for 20 min in a heptane-fixative mix at 3:1. The fixative consisted of 4% formaldehyde (Polysciences) in a PBS + 0.5% NP-40 (Sigma-Aldrich) solution. The ovarioles were partly detached, and ovaries were incubated for 2 hours in PBS-T (0.2% Tween 20) supplemented with 1% Triton X-100 (Sigma-Aldrich), 1% (w/v) bovine serum albumin and 1% (w/v) donkey serum. Primary antibody incubation was performed overnight at 4°C in BBT. The following day, ovaries were washed and incubated for 1 hour at room temperature in the secondary antibody solution (in BBT). Filamentous actin (f-Act) was stained with phalloidin-TRITC (Sigma-Aldrich) at 1:200 in PBS-T for 20 minutes. DNA was stained with SYTOX Green and the ovaries were mounted in fluorescence mounting medium. Fluorescence images were acquired with the previously described set-up (see “Embryo immunofluorescence”). Four independent experiments were conducted for each experimental condition and the analysis was performed in a blinded fashion.

For the bright-field imaging of whole ovaries, these were photographed after fixation using a camera-coupled Leica MZ12.5 stereomicroscope.

### Meiotic metaphase I arrest analysis

A protocol preventing egg activation was used to analyze the true metaphase I arrest configuration of the mature *Drosophila* female gamete [30,65]. Briefly, virgin females were aged, in the absence of males, for 4 days in standard medium supplemented with fresh yeast paste. Ovaries (10 pairs per sample per experiment) were quickly dissected in modified Robb’s medium and immediately transferred to fixative (4% formaldehyde in Robb’s medium). After a 5 minute incubation, ovaries were washed and DNA was stained with SYTOX Green. Two independent experiments were conducted for each experimental condition. Since this experiment (**Fig. 1F**) was part of a larger dataset, the quantification of the control group has already been published [30].

### Prophase I oocyte chromatin architecture analysis

The entire chromatin volume of individual prophase I oocytes was acquired as 0.5 μm-thick slices using a Leica SP5 confocal microscope (in fluorescence mounting medium). Two developmental stages were selected: before and after the establishment of the prophase I arrest (stages 1 and 6, respectively). Orb staining was used to unequivocally distinguish stage 1 oocytes from other neighbouring cells. Slices were stacked into maximum intensity Z-projections and binarized using an automated global thresholding method (Li’s minimum cross entropy thresholding; ImageJ). The perimeter of the binarized chromatin signal was then measured. Two independent experiments were conducted for each experimental condition and the analysis was performed in a blinded fashion.

### Embryo gene expression analysis

After manual isolation of pre-ZGA embryos (see “Protein immunoblotting”), total RNA was extracted using the PureLink RNA Mini Kit (Ambion). Total RNA was then processed for hybridization onto Drosophila Gene 1.1 ST Array Strip (Affymetrix) according to manufacturer’s instructions. For this, a GeneChip WT PLUS Reagent Kit (Affymetrix) and a GeneAtlas Hybridization, Wash and Stain Kit for WT Array Strips (Affymetrix) were used. One hundred nanograms of total RNA containing spiked in Poly-A RNA controls (GeneChip Poly-A RNA Control Kit, Affymetrix) were reverse transcribed for synthesis of single-stranded cDNA. After second-strand synthesis, the obtained double-stranded cDNA served as template for an *in vitro* transcription reaction to generate amplified cRNA. Fifteen micrograms of purified cRNA were then used for a second cycle of single-strand cDNA synthesis, after which 5.5 μg of ss-cDNA was fragmented and end-labeled. During this process, checkpoints for purity as well as integrity and size distribution of cRNA and fragmented ss-cDNA were performed using NanoDrop 1000 Spectrophotometer and Bioanalyzer 2100, respectively. Finally, 3.5 μg of the amplified, fragmented and labeled ss-cDNA were prepared in a 150 μl hybridization cocktail (containing hybridization controls) and 120 μl were used for the hybridization. The array strips were subsequently washed, double-stained and scanned. The hybridization, washing and scanning were performed using the Affymetrix GeneAtlas System.

The expression level of selected transcripts was validated by real-time quantitative reverse transcription PCR (qRT-PCR). Total RNA was extracted as before and 1 μg was used for reverse transcription with Oligo(dT**)18** primers (Transcriptor First Strand cDNA Synthesis Kit, Roche). qRT-PCR was performed under standard conditions using the Power SYBR Green PCR Master Mix (Applied Biosystems) in an ABI QuantStudio 7 station (Applied Biosystems). Samples were normalized using the expression of the Ribosomal protein L32 *(RpL32)* housekeeping gene. Relative expression levels were calculated with the 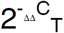 method, as previously described [66]. Primers were designed with Primer-BLAST (https://www.ncbi.nlm.nih.gov/tools/primer-blast/) and the corresponding sequences can be found in **Supplementary Table 4**.

For both the microarray and qRT-PCR analyses, two temporally independent experiments were conducted for each experimental condition.

### Embryo chromatin signal quantification

The entire volume of zygotic (cycle 1 or cycles 4 to 6) and polar body chromatin was acquired as 1 μm-thick slices using a Leica SP5 confocal microscope. Slices corresponding to each nucleus were stacked into a single maximum intensity Z-projection using ImageJ, and the limits of the chromatin area were defined. The mean fluorescence value of the tested signals was then recorded in arbitrary units (AU) inside the defined area, alongside that of a background reading. Relative signals were calculated as the mean fluorescence of the chromatin signal divided by the mean fluorescence of the corresponding background. A minimum of three independent experiments was conducted for each experimental condition.

### Statistical analysis

The scanned arrays were analyzed with Affymetrix Expression Console software using Robust Multi-array Analysis for both quality control and to obtain expression values. Control probe sets were removed and log2 expression values of the remaining 15308 transcripts were imported into Chipster 3.8.1 [67]. Differential expression was determined by empirical Bayes two-group test with Benjamini-Hochberg multiple testing correction and a p-value cut-off of 0.05 [68].

Gene Ontology analysis was performed using the Metascape Gene Annotation and Analysis Resource tool (http://metascape.org/gp/index.html#/main/step1) using the “express analysis” settings [69].

Nonparametric tests (Mann-Whitney U test) were used to compare egg size, relative fluorescence signals and chromatin perimeter measurements between groups. For egg eclosion and mitotic entry rates, unpaired (two sample) t-tests were used. The comparison of male PN configuration, sperm ProtB-GFP signal, meiotic completion, MI arrest configuration and early zygotic development between groups was performed by two-way ANOVA. Reported *P* values correspond to two-tailed tests. All analyses were performed using Prism 7 software (GraphPad).

### Data availability

All relevant data and reagents are available from the Authors. The microarray data discussed in this publication have been deposited in NCBI’s Gene Expression Omnibus [70] and are accessible through GEO Series accession number GSE108033 (https://www.ncbi.nlm.nih.gov/geo/query/acc.cgi?acc=GSE108033).

## ACKNOWLEDGEMENTS

The authors wish to thank Gabriel Martins and Nuno Pimpão for assistance in fluorescence microscopy. We acknowledge Renate Renkawitz-Pohl (Philipps-Universität Marburg, Germany) and Mariana Wolfner (Cornell University, Ithaca, USA) for kindly providing us with the ProtB-GFP and *sarah* mutant fly stocks, respectively. We also thank Alexander Mazo (Thomas Jefferson University, Philadelphia, USA) and Ali Shilatifard (Northwestern University Feinberg School of Medicine, Chicago, USA) for aliquots of an anti-dMLL3/4 antibody. We acknowledge the TRiP at Harvard Medical School (NIH/NIGMS R01-GM084947) for providing several of the transgenic RNAi fly stocks used in this study. Rui G. Martinho is supported by Portuguese national funding through the following Fundação para a Ciência e a Tecnologia (FCT) grants: PTDC/BEX-BID/0395/2014 and UID/BIM/04773/2013 CBMR 1334. Jörg D. Becker received salary support from FCT through an “Investigador FCT” position (IF/01341/2012). Paulo Navarro-Costa is supported by a FCT Postdoctoral fellowship (SFRH/BPD/84214/2012).

## AUTHORS CONTRIBUTIONS

P.P.: Investigation. L.G.G.: Investigation. J.S.: Investigation. J.D.B.: Investigation, Writing (review & editing). R.G.M: Conceptualization, Writing (review & editing), Funding acquisition. P.N-C: Conceptualization, Investigation, Writing (original draft + review & editing).

